# Single-dose Efficacy of a Next-Generation Mpox Vaccine Harnessing an Immunomodulatory Peptide

**DOI:** 10.64898/2026.07.31.742137

**Authors:** Anoli Karunathilake, Reem Miah, Crystal Lawson, JD Burleson, Sreenivasa RaoOruganti, Mary J Hauser, Arban Domi, Brooklyn Adragna, Brayden Olsen, Shuangyi Bai, Pratima Kumari, Bronwyn M Gunn, Mark Newman, Heather Koehler

## Abstract

Rapidly deployable, single-dose vaccines that maintain durability under operational constraints remain an unmet need in outbreak preparedness. Live viral vectors such as Modified Vaccinia Ankara (MVA) offer strong safety profiles, yet their suboptimal immunogenicity often requires multidose regimens, reducing flexibility during emergency response. To address these limitations, we developed a modular vaccine platform that leverages immune checkpoint modulation to enhance immune cell priming without compromising the established safety profile of MVA. This platform, exemplified by the recombinant virus MVA-X, was engineered to express a peptide-based PD-1 antagonist (LD10) that provides localized, transient checkpoint blockade during early antigen presentation. The approach requires no external adjuvants, is compatible with lyophilization and stockpiling, and is readily adaptable to diverse antigens and pathogens.

A single immunization with MVA-X produced durable protection that matched or exceeded that of a conventional two-dose MVA regimen against the prototypic orthopoxvirus vaccinia virus. Despite modest and contracting antibody titers, single-dose MVA-X vaccination conferred complete survival following both lethal and high-dose viral challenge at early (Day 55), intermediate (Day 90), and long-term (Day 150) time points. MVA-X also restricted viral replication at the primary site of infection, reduced systemic dissemination, and preserved lung architecture during peak disease. Importantly, MVA-X maintained efficacy in the highly susceptible CAST/EiJ mouse model following challenge with highly pathogenic Clade I monkeypox virus (MPXV). Together, these findings demonstrate that vaccine-intrinsic checkpoint modulation provides a modular strategy for enhancing the potency and durability of attenuated viral vectors while preserving their favorable safety profile, supporting broader application to emerging infectious diseases beyond Mpox.

## Introduction

The development of vaccines that are both exceptionally safe and capable of inducing durable protection after a single administration remains a central challenge in outbreak preparedness^1–3^. Although many licensed vaccines rely on booster dosing to achieve sustained immunity, follow-up vaccination can be difficult during public health emergencies because of limited infrastructure, disrupted supply chains, and reduced access to healthcare^4–6^. Consequently, vaccine strategies capable of inducing durable protection after a single immunization remain a challenge.

Modified Vaccinia Ankara (MVA) is a live, highly attenuated poxvirus vector with an extensive safety record across diverse populations, including immunocompromised individuals ^7–11^. Its large genome capacity, genetic stability, and compatibility with scalable manufacturing have led to its widespread development as a vaccine platform for infectious diseases and cancer^12–14^. MVA-based vaccines are also amenable to stockpiling and lyophilization, features that facilitate rapid deployment during outbreak response^15^. However, the same attenuation that underlies its favorable safety profile can limit the magnitude and durability of immune responses following a single immunization, often necessitating booster doses to achieve optimal protection^16^. Although a two-dose MVA-BN regimen remains the recommended standard for Mpox vaccination, continued evaluation of additional booster regimens highlights ongoing efforts to further optimize the magnitude and durability of MVA-induced immunity^17^. These approaches primarily seek to enhance protection by increasing vaccine exposure, whereas strategies that improve the quality of immune priming after a single immunization remain comparatively underexplored. This trade-off between safety and immunogenicity constrains the operational flexibility of MVA vaccination strategies, particularly when rapid population-level coverage is required^18^. These limitations became evident during the 2022 global Mpox outbreak, when MVA-based vaccines were deployed under emergency use conditions^19^. Although vaccination reduced severe disease and hospitalization, real-world studies demonstrated incomplete protection following a single dose, delayed booster uptake, and challenges associated with vaccine availability and distribution^6,20–22^.

Protective immunity against orthopoxvirus infection depends on coordinated humoral and cellular immune responses^23^. Neutralizing antibodies contributes to limiting viral dissemination, whereas CD8⁺ T cells are essential for eliminating infected cells and establishing durable protection against disease^9,24^. Studies of both Vaccinia virus and mpox virus infection have shown that robust cellular immunity is associated with improved viral control, reduced disease severity, and enhanced survival, particularly as circulating antibody responses over time^25^. In addition to direct cytolytic activity, antigen-specific CD8⁺ T cells produce antiviral cytokines, including IFNγ and TNFα, that suppress viral replication and amplify local antiviral responses^26–28^.

However, real-world experience during recent Mpox outbreaks, together with emerging longitudinal immunogenicity studies, indicates that although MVA-BN induces both humoral and cellular immune responses, durable immunity following a single immunization remains suboptimal. As a replication-deficient vector, MVA provides an excellent safety profile but reduced antigen persistence and inflammatory signaling compared with replication-competent orthopoxviruses vaccines^29,30^, potentially limiting the magnitude and durability of adaptive immune responses after a single dose. Consistent with this biology, both binding antibody and antigen-specific T-cell responses progressively contract over time, with cellular immunity persisting longer than humoral immunity but nonetheless substantially diminished during extended follow-up^30–33^. Collectively, these findings identify enhancement of durable immune priming as a promising strategy for improving single-dose MVA vaccination while preserving its established safety profile.

These observations suggest that enhancing the quality of vaccine-induced cellular immunity may be an effective strategy for improving the durability of single-dose MVA vaccination. The programmed cell death protein-1 (PD-1) pathway regulates T-cell activation, differentiation, and memory formation during antigen priming^34–36^. Transient modulation of PD-1 signaling at the time of vaccination has been shown to enhance effector expansion and memory development in preclinical models^37^. However, systemic or prolonged checkpoint inhibition is not well suited for prophylactic vaccination because of the potential for immune-related adverse events^38^. An approach that enables localized and self-limited checkpoint modulation during antigen presentation could therefore enhance immune priming while preserving safety. Multiple preclinical studies have demonstrated that transient modulation of immune checkpoint pathways during vaccination enhances antigen-specific CD8⁺ T-cell expansion, effector function, and memory formation, including when PD-1- and CTLA-4-targeted strategies are incorporated into viral-vectored vaccines such as adenoviral and poxviral platforms^39–41^.

Here, we developed MVA-X, a recombinant MVA vaccine engineered to incorporate transient immune checkpoint modulation directly within the viral vector. MVA-X expresses LD10, a genetically encoded 18-amino-acid peptide-based PD-1 antagonist that was identified through microbiome mining of the *Bacillus thuringiensis* Cry1A toxin and shares sequence similarity with a previously characterized PD-1 antagonist^42,43^. By encoding immune modulation directly within the vaccine vector, this strategy is designed to enhance T-cell priming without requiring external adjuvants or systemic checkpoint inhibition.

Unlike conventional checkpoint inhibitors, MVA-X was specifically designed to confine PD-1 antagonism to the site and timing of vaccine-induced immune priming. MVA preferentially infects professional antigen-presenting cells, particularly dendritic cells responsible for initiating adaptive immune responses^44^, thereby localizing LD10 expression to the site of antigen presentation. Because MVA is replication-deficient in mammalian cells and is rapidly cleared following vaccination^45^, transgene expression is restricted to the early stages of the immune response. Furthermore, LD10 exhibits a short in vivo half-life of approximately one hour following administration in mice, further limiting checkpoint modulation^37^. Together, these characteristics are expected to restrict PD-1 antagonism both spatially and temporally, enhancing early T-cell priming while minimizing prolonged systemic immune activation, an approach previously validated in a malaria vaccine model^37^.

We evaluated MVA-X in murine models of lethal intranasal vaccinia virus Western Reserve (VACV-WR) challenge to determine whether vector-intrinsic checkpoint modulation could improve single-dose protection and durability. Lethal VACV challenge provides a rigorous model for assessing the magnitude and longevity of orthopoxvirus immunity under conditions of high viral burden. Protection was evaluated early (Day 55), intermediate (Day 90), and long-term (Day 150) time points to determine whether durable immunity could be maintained despite declining antibody responses. Because protection against Mpox remains a major public health priority, vaccine efficacy was further assessed in the highly susceptible CAST/EiJ mouse model following challenge with highly pathogenic Clade I MPXV, providing a stringent evaluation of vaccine performance against severe orthopoxvirus disease. Collectively, these studies examine whether localized immune checkpoint modulation can improve the durability of established viral-vector vaccines without compromising their underlying safety profile.

## Results

### Construction and validation of the checkpoint-integrated MVA-X vaccine platform

To enhance T-cell priming within the established safety framework of Modified Vaccinia Ankara (MVA), we engineered a recombinant MVA vector expressing the peptide-based PD-1 antagonist LD10, designated MVA-X. The LD10 expression cassette was inserted between two essential viral genes under the control of an MVA-specific promoter and consisted of five tandem LD10 repeats, each preceded by a secretion signal peptide and separated by proteolytic cleavage motifs to facilitate peptide processing and extracellular release (Fig. 1). Expression and secretion of LD10 from infected cells were confirmed by dot blot analysis of culture supernatants (Supplementary Fig. 1), consistent with previous characterization of the construct promoter (Fig. 1)^37^.

**Figure 1:**
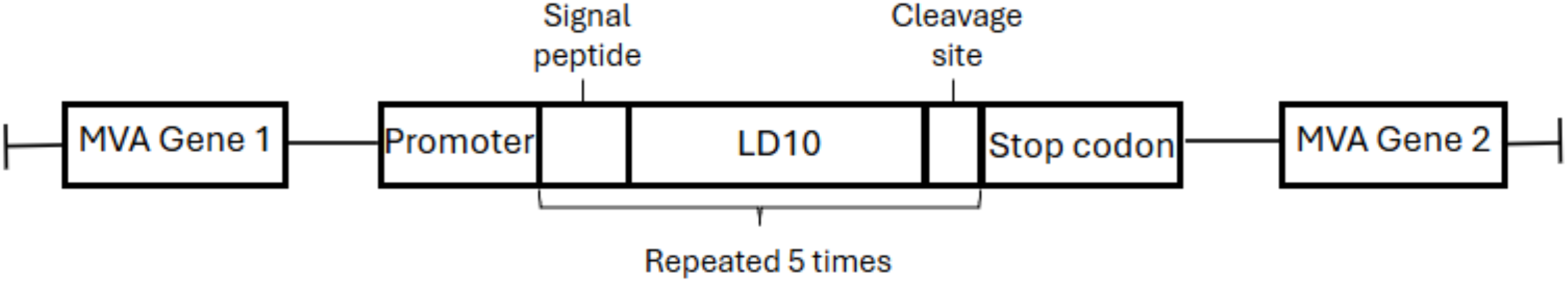
Schematic of recombinant MVA-X construct. The LD10 sequence inserted into the MVA genome between two essential genes under control of an MVA specific promoter. LD10 sequences are preceded by a signal sequence routing peptides for secretion and followed by a cleavage site to separate duplicated peptides. The secretion signal, LD sequence and cleavage site are repeated 5 times and then transcription is terminated with a stop codon.

### MVA-specific binding and neutralizing antibody responses contract by D90 post-prime vaccination

To characterize vaccine-induced humoral immunity, MVA-specific serum IgG binding antibodies were measured longitudinally following vaccination. At Day 55 post-prime vaccination, mice receiving two-dose MVA (MVA Prime/Boost) exhibited the highest endpoint IgG titers, whereas single-dose MVA (MVA Prime) and MVA-X (MVA-X Prime) elicited lower responses, and mock-vaccinated animals remained seronegative (Fig. 2A). Binding antibody titers declined by Days 75 and 90 across all vaccinated groups, with the greatest reduction observed in the MVA Prime/Boost group.

**Figure 2:**
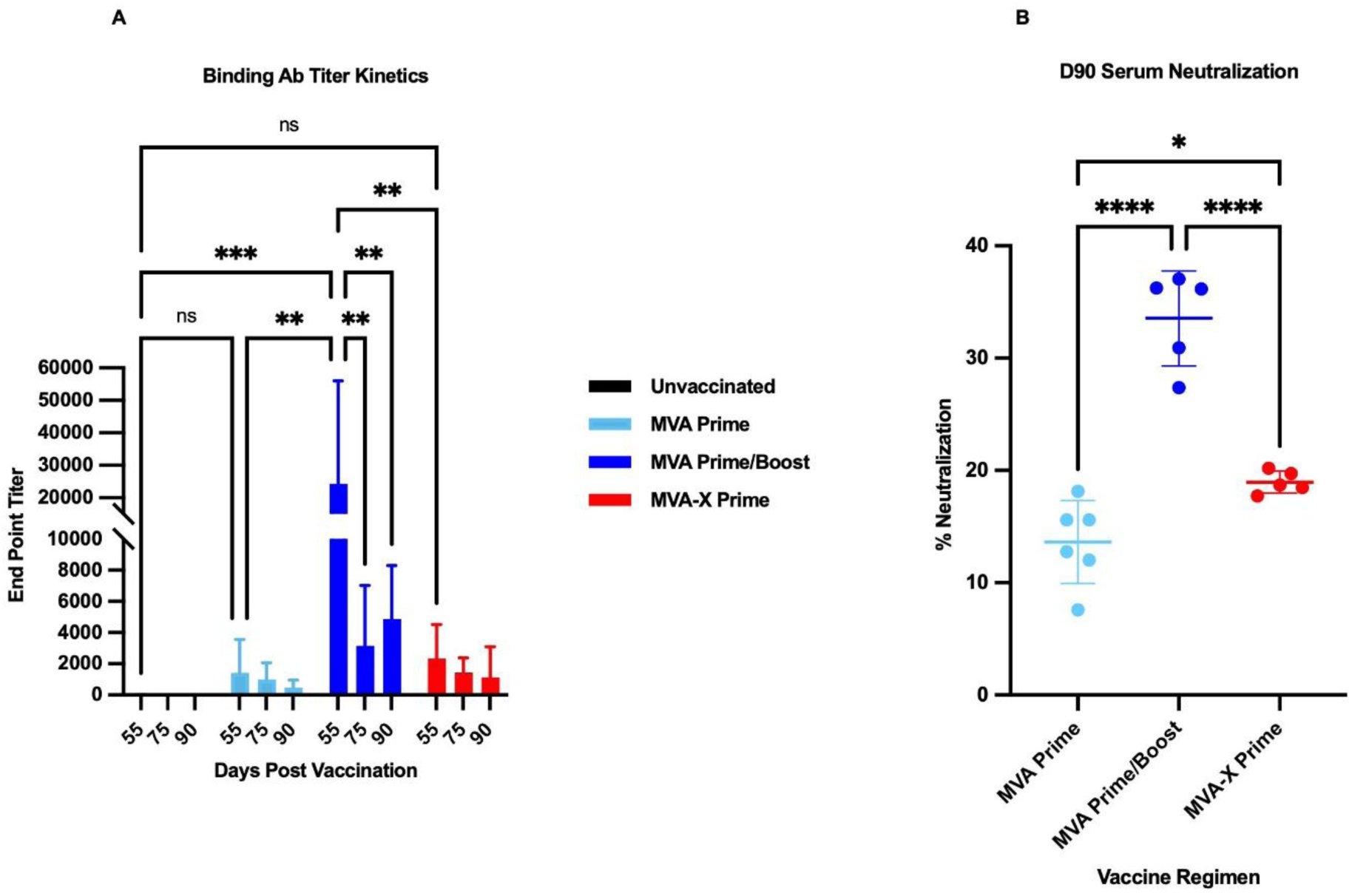
MVA-specific binding and neutralizing antibody responses contract by Day 90 post-prime vaccination. **(A)** Longitudinal MVA-specific serum IgG endpoint titers measured by ELISA at Days 55, 75 and 90 post-prime vaccination. Groups include unvaccinated mice, MVA Prime, MVA Prime/Boost, and MVA-X Prime. Endpoint titers were defined as the highest serum dilution producing absorbance values at least two-fold above background. Each bar represents the group mean with standard deviation (n = 5–6). On Day 55, the MVA Prime/ Boost group showed significantly higher binding antibody titers than unvaccinated, MVA Prime, and MVA- X Prime groups. Binding titers in the MVA Prime/Boost group declined significantly between Day 55 and Days 75 and 90, whereas titers in the remaining groups remained low and were not significantly different across time. A declining trend was observed across vaccination regimens. (N= 5-6)**. (B)** Serum neutralizing activity measured at Day 90 post-prime using a VACV-E3L- GFP flow cytometry-based neutralization assay. Neutralization is expressed as the percentage reduction in GFP-positive infected Vero-81 cells relative to virus-only controls at the same multiplicity of infection. Individual points represent individual mice; horizontal bars indicate mean with standard deviation. Two-dose MVA Prime/Boost group induced the highest neutralizing activity and was significantly greater than both MVA Prime and MVA-X Prime. MVA-X Prime showed a modest but significant increase in neutralization compared with MVA Prime. Statistical analysis was performed using two-way ANOVA with Tukey’s multiple-comparisons test for panel 2A and ordinary one-way ANOVA with Tukey’s multiple-comparisons test for panel 2B. Exact statistical comparisons are indicated in the figure. ns, not significant; *P < 0.05; **P < 0.01; ***P < 0.001; ****P < 0.0001.

Serum neutralizing activity was assessed at Day 90 using a flow cytometry-based VACV-E3L- GFP neutralization assay. Consistent with the binding antibody responses, the MVA Prime/Boost group exhibited the highest neutralizing activity (Fig. 2B). Single-dose MVA-X induced modest but detectable neutralizing responses that were significantly greater than those elicited by parental MVA Prime alone but remained lower than the Prime/Boost regimen, consistent with previous observations of MVA-induced neutralizing antibody response studies (Fig. 2B) ^46^.

### Single-dose MVA-X confers robust protection against lethal VACV-WR challenge at Days 55 and 90 post-prime vaccination

To determine whether vaccine-induced humoral responses translated into protective immunity, mice were challenged intranasally with a lethal dose (1 × 10⁷ PFU) of VACV-WR at either Day 55 or Day 90 post-prime vaccination and monitored for morbidity and survival.

Following Day 55 challenge, unvaccinated animals developed progressive weight loss beginning at approximately 3 days post-infection (dpi) and succumbed uniformly by 6 dpi (Fig. 3A, B). In contrast, both MVA-X Prime and MVA Prime/Boost vaccination conferred complete protection, maintaining body weight and 100% survival throughout the observation period. Mice receiving a single dose of parental MVA (MVA Prime) exhibited transient weight loss and partial protection, with one mortality occurring during infection.

**Figure 3:**
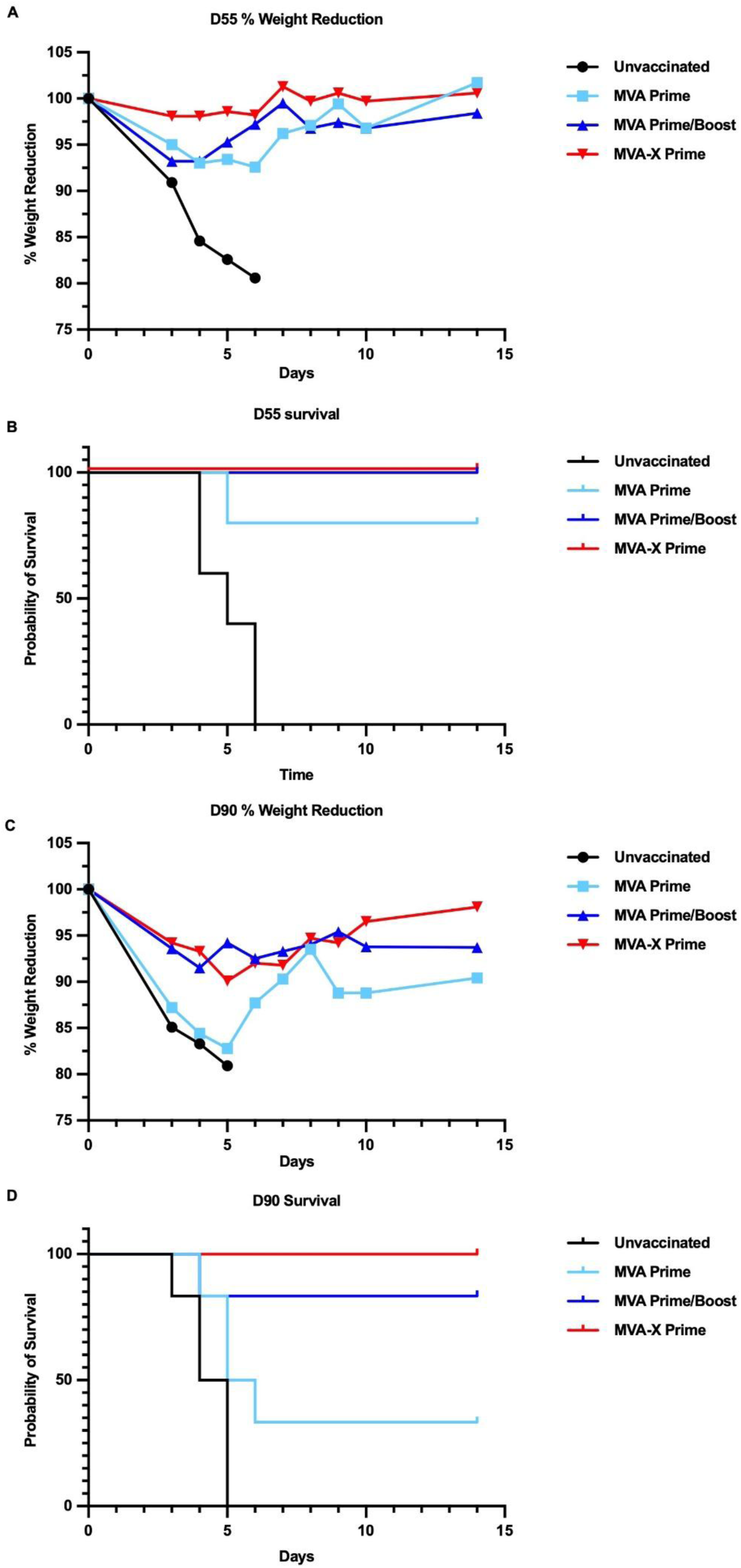
Single-dose MVA-X confers robust protection against lethal VACV-WR challenge at Days 55 and 90 post-prime vaccination. (**A-B**) Mice (n = 6 per group) were challenged intranasally with 1 × 10⁷ PFU VACV-WR at Day 55 post-prime vaccination and monitored for 14 d post- infection (dpi). Group averages are shown for body weight measurements. (**A**) Percent body weight relative to pre-challenge baseline over the course of infection. Unvaccinated animals developed rapid and progressive weight loss beginning at approximately 3 dpi. In contrast, mice receiving MVA-X Prime or MVA Prime/Boost maintained stable body weights throughout the observation period, whereas Single-dose parental MVA vaccination resulted in transient weight loss followed by recovery. (**B**) Kaplan–Meier survival analysis following Day 55 challenge All unvaccinated animals succumbed to infection by 6 dpi, with deaths occurring at 4 dpi (n = 2), 5 dpi (n = 1), and 6 dpi (n = 3). A single mortality was observed in the parental MVA Prime group at 4 dpi, whereas complete survival was maintained in both the MVA-X Prime and MVA Prime/Boost groups throughout the study period. **(C-D)** Mice (n = 6 per group) were challenged intranasally with 1 × 10⁷ PFU VACV-WR at Day 90 post-prime vaccination and monitored for 14 dpi. Group averages are shown for body weight measurements. (**C**) Percent body weight relative to pre-challenge baseline following infection. Unvaccinated mice exhibited severe disease progression with marked weight loss, while MVA-X Prime vaccination preserved body weight stability throughout infection. The MVA Prime and MVA Prime/Boost groups showed intermediate transient weight loss with partial recovery. (**D**) Kaplan–Meier survival analysis following Day 90 challenge. All unvaccinated animals succumbed by 5 dpi, with deaths occurring at 4 dpi (n = 3) and 5 dpi (n = 3). In the parental MVA Prime group, deaths occurred at 4 dpi (n = 1), 5 dpi (n = 1), and 6 dpi (n = 1), resulting in 50% survival. A single mortality occurred in the MVA Prime/Boost group at 5 dpi, whereas complete protection was maintained in the MVA-X Prime group throughout the 14-day observation period. Survival curves were analyzed using the log-rank (Mantel–Cox) test. Group comparisons and exact statistical analyses are indicated in the figure.

Protection conferred by MVA-X was maintained on Day 90 despite declining humoral responses. Unvaccinated animals again developed severe disease characterized by rapid weight loss and complete mortality by 5 dpi (Fig. 3C, D). Single-dose MVA-X vaccination preserved body weight and maintained 100% survival, whereas MVA Prime and MVA Prime/Boost groups exhibited greater morbidity with reduced survival (50% and 83%, respectively).

### MVA-X suppresses pulmonary viral burden and limits systemic dissemination following lethal VACV-WR challenge

To determine whether the enhanced protection conferred by MVA-X vaccination was associated with improved control of viral replication, viral genome burden was quantified in lung and blood samples collected at 5 dpi following Day 90 lethal VACV-WR challenge.

Quantitative PCR targeting the conserved VACV **A12L** gene demonstrated high pulmonary viral genome levels in unvaccinated animals and mice receiving a single dose of parental MVA (Fig. 4A). In contrast, both MVA-X Prime and MVA Prime/Boost vaccination significantly reduced viral genome levels within the lung, indicating effective restriction of viral replication at the primary site of infection.

**Figure 4.**
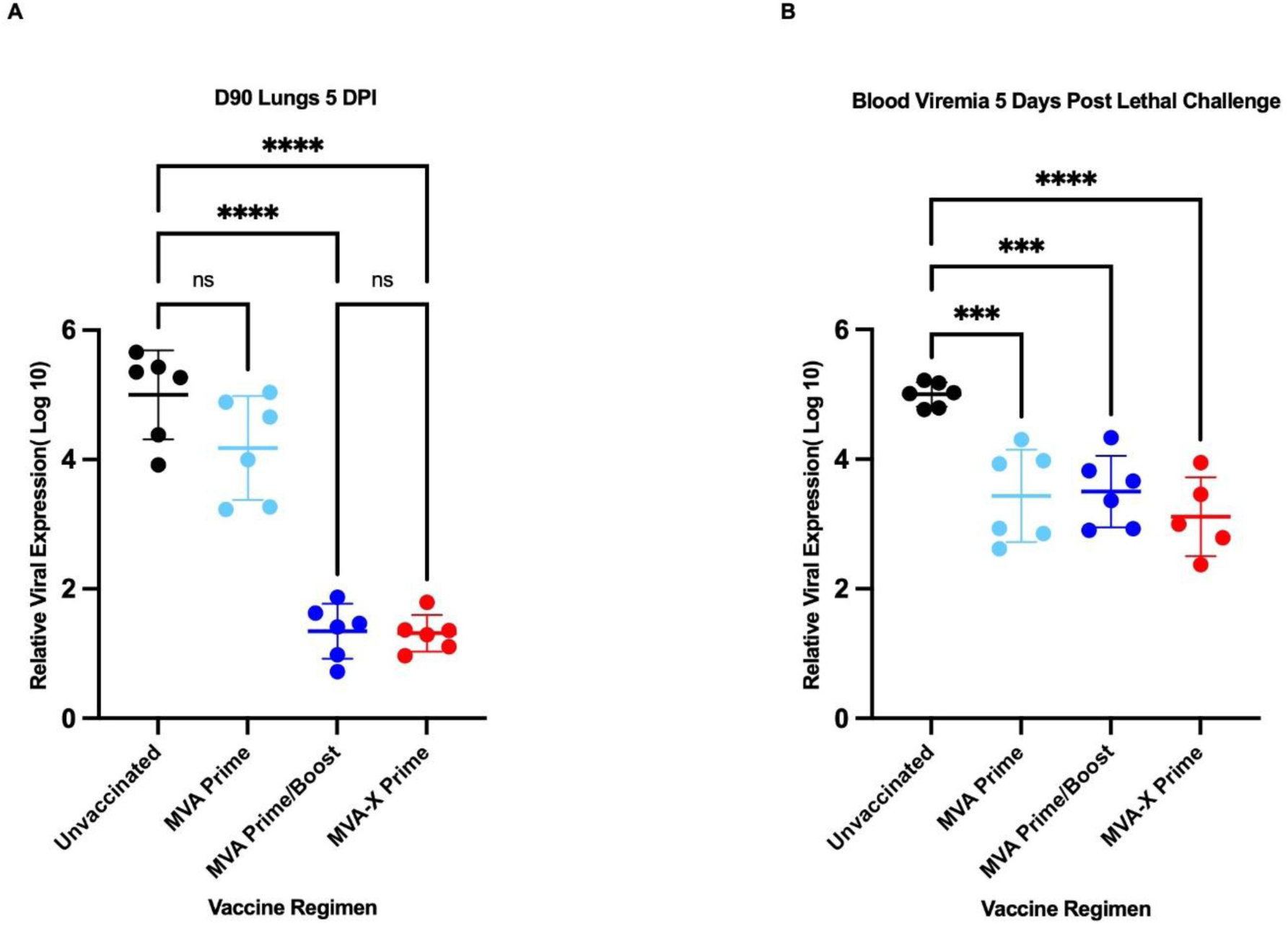
MVA-X suppresses pulmonary viral burden and limits systemic dissemination following lethal VACV-WR challenge. **(A)** Relative pulmonary viral genome levels measured in lung tissue collected at 5 d post-infection (dpi) following Day 90 intranasal challenge with 1 × 10⁷ PFU VACV-WR. Viral DNA was quantified by qPCR targeting the VACV A12L gene and normalized to β-actin expression. Relative viral expression is presented as log₁₀(2^−ΔΔCt) + 5, where the unvaccinated group served as the reference comparator for ΔΔCt calculations and 5 was added as a visualization constant. Individual points represent individual animals (n = 6 per group); horizontal bars indicate mean ± s.d. Unvaccinated and Single-dose MVA Prime groups exhibited high pulmonary viral burden, whereas both MVA Prime/Boost and MVA-X Prime groups showed marked suppression of viral genomes within the lung**. (B)** Relative blood viral genome levels measured at 5 dpi following Day 90 VACV-WR challenge. Viral DNA quantification, normalization, and transformation were performed as described in panel a. Individual points represent individual animals (n = 5–6 per group); horizontal bars indicate mean ± s.d. All vaccinated groups demonstrated reduced systemic viral burden relative to unvaccinated controls, with the greatest reduction observed in the MVA-X Prime group. Statistical analysis was performed using ordinary one-way ANOVA with Tukey’s multiple- comparisons test. Exact statistical comparisons are indicated in the figure. ns, not significant; ***P < 0.001; ***P < 0.0001

Analysis of circulating viral genomes revealed a similar trend (Fig. 4B). All vaccinated groups exhibited reduced blood viral genome levels relative to unvaccinated controls, with the greatest reduction observed in the MVA-X Prime group. These findings demonstrate that a single administration of MVA-X limits both local pulmonary viral replication and systemic viral dissemination following high-dose orthopoxvirus challenge.

### Single-dose MVA-X restricts pulmonary viral replication and preserves lung architecture following lethal VACV-WR challenge

To determine whether reduced viral genome burden translated into effective control of infectious virus and preservation of tissue integrity, infectious viral titers and lung histopathology were evaluated at 5 dpi following Day 90 lethal VACV-WR challenge.

Unvaccinated animals exhibited high pulmonary viral titers, consistent with uncontrolled viral replication at the primary site of infection (Fig. 5A). In contrast, both MVA-X Prime and MVA Prime/Boost vaccination markedly reduced infectious viral burden within the lung. Mice receiving a single dose of parental MVA demonstrated intermediate viral control with substantial inter-animal variability.

**Figure 5:**
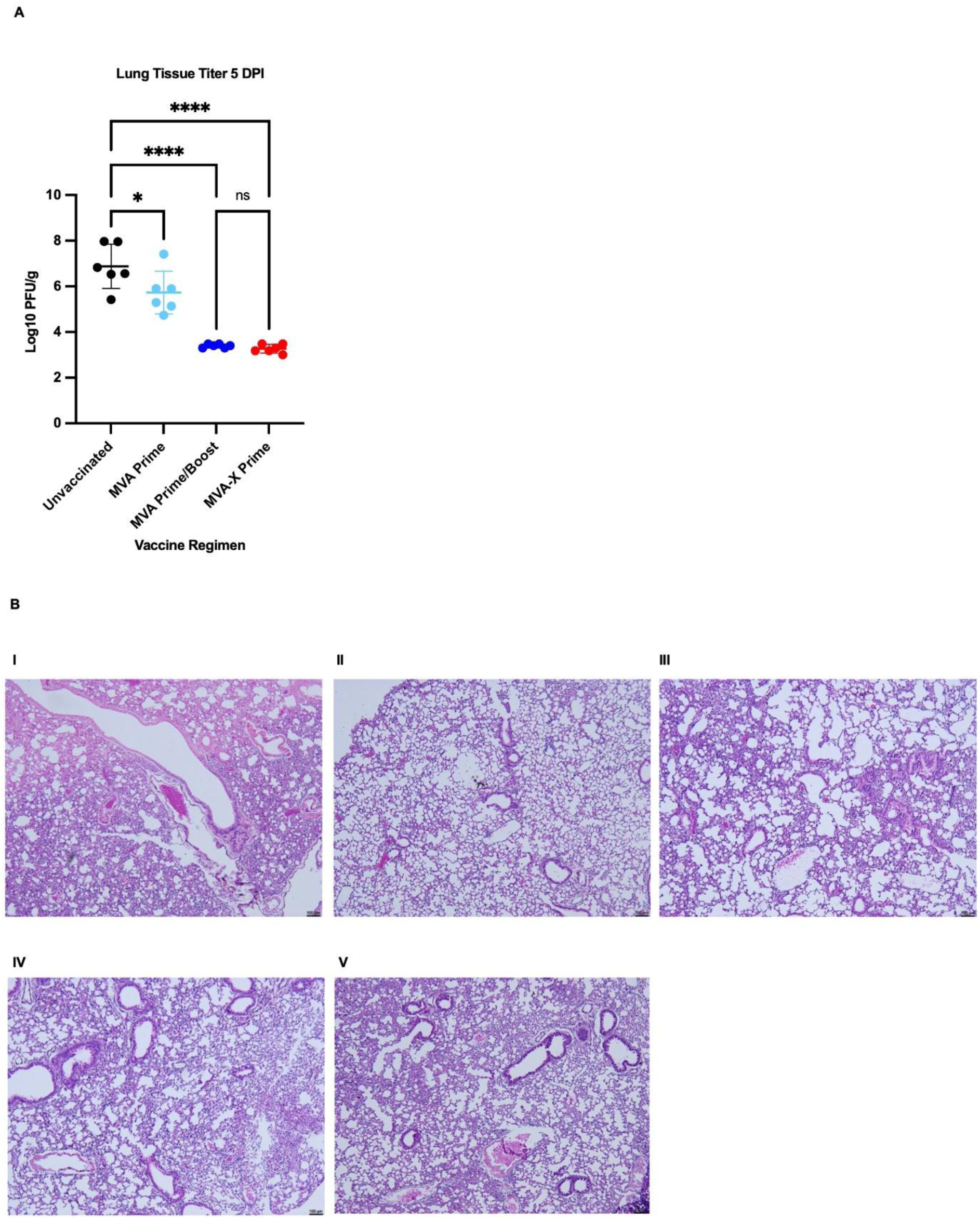
MVA-X Prime restricts pulmonary viral replication and preserves lung architecture following lethal VACV-WR challenge. **(A)** Lung viral titers measured at 5 d post-infection (dpi) following Day 90 intranasal challenge with 1 × 10⁷ PFU VACV-WR. Infectious viral burden was quantified by plaque assay on Vero-81 cells and is presented as log₁₀ plaque- forming units (PFU) per gram of lung tissue. Individual points represent individual animals (n = 6 per group); horizontal bars indicate mean ± s.d. Unvaccinated animals exhibited high pulmonary viral titers, while both MVA Prime/Boost and MVA-X Prime groups showed marked suppression of infectious viral replication within the lung. The Single-dose parental MVA group demonstrated intermediate viral control with greater inter-animal variability. **(B)** Representative hematoxylin and eosin (H&E)-stained lung sections collected at 5 dpi following Day 90 VACV-WR challenge. Images are shown at 5× magnification. Healthy uninfected lungs displayed intact alveolar architecture with open airspaces and no evidence of inflammation (I). Mock-vaccinated animals exhibited extensive alveolar destruction, inflammatory consolidation, and severe pulmonary pathology consistent with uncontrolled viral infection (II). Lungs from mice receiving Single-dose parental MVA showed partial preservation of tissue architecture with localized inflammatory infiltrates and areas of alveolar collapse (III). In contrast, homologous MVA Prime/Boost and Single-dose MVA-X vaccination preserved overall lung structure with organized immune infiltration and reduced tissue destruction (IV, V). Scale bar, 100 μm. Statistical analysis for panel a was performed using ordinary one-way ANOVA with Tukey’s multiple-comparisons test. Exact statistical comparisons are indicated in the figure. ns, not significant; *P < 0.05; ****P < 0.0001.

Histopathological analysis revealed marked differences in disease severity between vaccination groups (Fig. 5B). Unvaccinated animals developed extensive pulmonary pathology characterized by alveolar destruction, inflammatory consolidation, and widespread tissue disruption. MVA Prime vaccination resulted in partial preservation of lung architecture, although inflammatory infiltrates and alveolar collapse remained evident. In contrast, lungs from MVA-X Prime-vaccinated animals retained preserved alveolar architecture with organized immune cell infiltration, comparable to that observed in the MVA Prime/Boost group. Together, these findings demonstrate that MVA-X vaccination effectively limits pulmonary viral replication while preserving lung architecture following lethal orthopoxvirus challenge.

### MVA-X enhances durable antigen-specific CD8⁺ T cell immunity associated with protection following lethal VACV-WR challenge

To determine whether the durable protection conferred by MVA-X was associated with enhanced cellular immunity, antigen-specific CD8⁺ T-cell responses were evaluated at Day 90 post-prime vaccination. Day 90 was selected because it represents a post-contraction memory time point at which circulating antibody responses had declined while protection against lethal VACV challenge remained intact. Splenocytes were stimulated ex vivo with whole MVA and analyzed by intracellular cytokine staining.

MVA-X Prime vaccination induced increased frequencies of IFNγ⁺ CD8⁺ T cells compared with unvaccinated and MVA Prime groups (Fig. 6A). Similarly, TNFα-producing CD8⁺ T-cell responses were elevated in the MVA-X Prime group relative to the remaining vaccination groups (Fig. 6B). In contrast, IL-2⁺ CD8⁺ T-cell frequencies showed limited separation between groups (Fig. 6C), suggesting preferential enhancement of effector-associated cytokine responses rather than broad expansion of all functional CD8⁺ T-cell subsets. To further determine whether MVA-X vaccination influenced the differentiation of memory CD8⁺ T-cell populations, effector memory (TEM) and central memory (TCM) subsets were evaluated at Day 90 post-prime vaccination. MVA-X Prime vaccination resulted in increased frequencies of CD8⁺ TEM cells compared with unvaccinated, single-dose MVA Prime, and MVA Prime/Boost groups (Supplementary Fig. 4), whereas TCM frequencies were not significantly different among vaccination regimens. These findings suggest that vaccine-intrinsic PD-1 modulation preferentially promotes the development or maintenance of effector memory CD8⁺ T cells, consistent with the durable protection observed following MVA-X vaccination.

**Figure 6:**
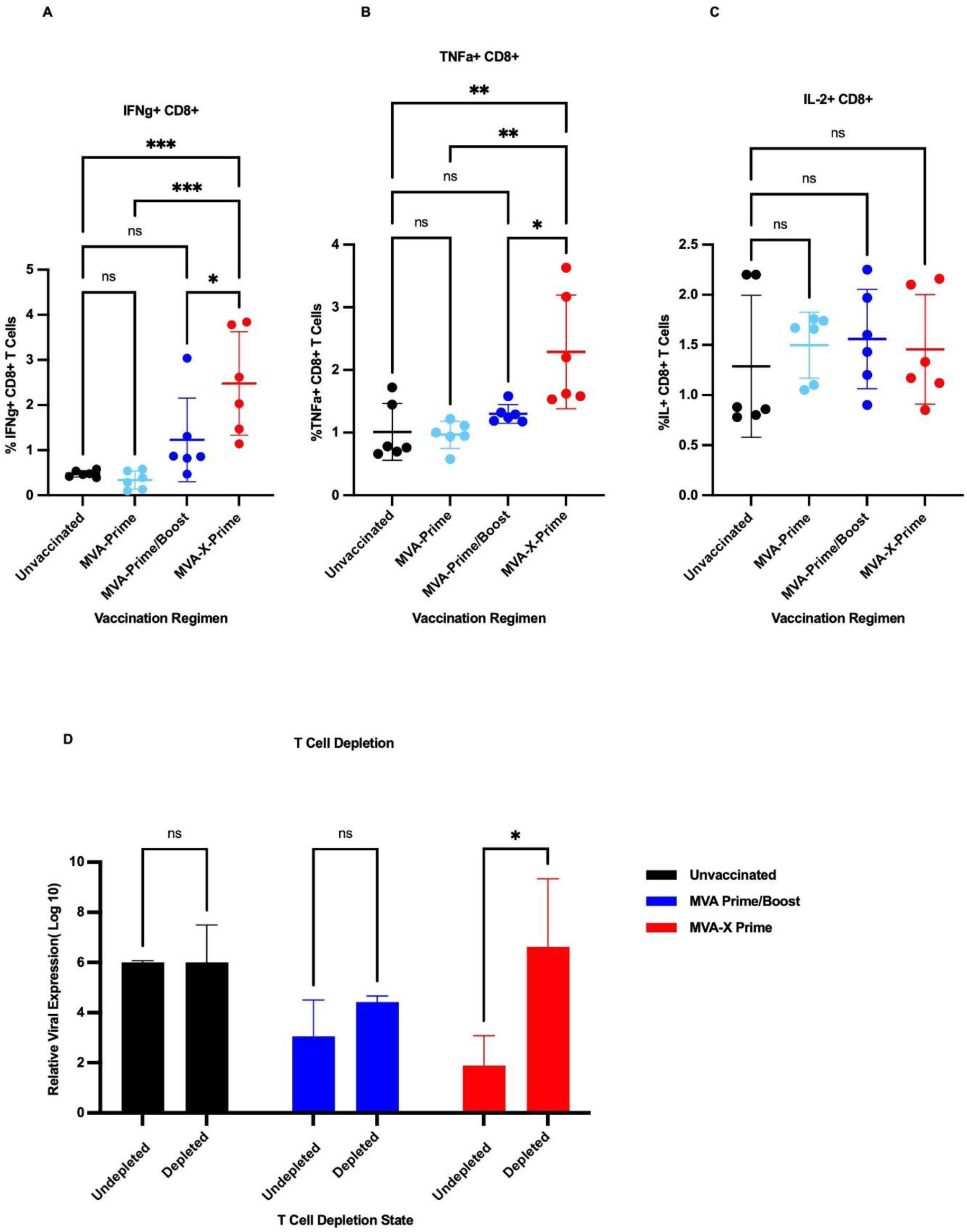
MVA-X Prime vaccination promotes durable antigen-specific CD8⁺ T cell immunity associated with enhanced viral control following lethal VACV-WR challenge. Splenocytes or tissues were collected from mice vaccinated with Single-dose MVA-X, Single-dose parental MVA, or two-dose parental MVA Prime/Boost regimens and analyzed at Day 90 post-prime vaccination. Individual points represent individual animals unless otherwise indicated; horizontal bars represent mean ± s.d. **(A)** Percentage of IFNγ⁺ CD8⁺ T cells following ex vivo stimulation of splenocytes with whole MVA and intracellular cytokine staining. MVA-X Prime vaccination generated elevated IFNγ-producing CD8⁺ T cell responses relative to parental MVA vaccination groups. **(B)** Percentage of TNFα⁺ CD8⁺ T cells following ex vivo MVA stimulation. Increased TNFα-producing CD8⁺ T cell responses were observed in the MVA-X Prime group compared with the remaining vaccination cohorts. **(C)** Percentage of IL-2⁺ CD8⁺ T cells following ex vivo MVA stimulation. IL-2-producing CD8⁺ T cell responses were detected across all vaccination groups with relatively limited intergroup variation. **(D)** Relative viral genome levels following CD8⁺ T cell depletion prior to VACV-WR challenge. Mice were treated with CD8⁺ T cell-depleting antibody before challenge, and viral genome burden was quantified by qPCR targeting the VACV A12L gene. Relative viral expression is presented as log₁₀(2^−ΔΔCt) + 5, with the undepleted unvaccinated group used as the reference comparator. CD8⁺ T cell depletion resulted in increased viral burden in the MVA-X Prime group, supporting a functional contribution of CD8⁺ T cell-mediated immunity to protection following MVA-X vaccination. Statistical analysis was performed using ordinary one-way ANOVA with Tukey’s multiple-comparisons test for panels a–c and two-way ANOVA with Tukey’s multiple- comparisons test for panel d. Exact statistical comparisons are indicated in the figure. ns, not significant; *P < 0.05; **P < 0.01; ***P < 0.001.

To determine whether these responses contributed functionally to protection, CD8⁺ T cells were depleted prior to Day 90 VACV-WR challenge. Efficient depletion was confirmed in peripheral blood before infection (Supplementary Fig. 5). CD8⁺ T-cell depletion had minimal effect on viral genome burden in unvaccinated animals or in the MVA Prime/Boost group. In contrast, depletion of CD8⁺ T cells in MVA-X-vaccinated animals resulted in a marked increase in viral genome burden at 5 dpi, approaching levels observed in unvaccinated controls (Fig. 6D). To assess whether innate cytotoxic lymphocytes also contributed to protection, NK cell depletion studies were performed. In contrast to CD8⁺ T-cell depletion, NK cell depletion did not significantly alter pulmonary viral burden following VACV challenge (Supplementary Fig. 6). These findings demonstrate that the enhanced protection conferred by single-dose MVA-X vaccination is substantially dependent on CD8⁺ T-cell immunity and supports a central role for vaccine-induced cellular responses in controlling viral replication following lethal orthopoxvirus challenge.

### Single-dose MVA-X maintains protection and antigen-specific CD8⁺ T-cell responses through Day 150 post-vaccination

To evaluate the long-term durability of MVA-X-induced immunity, mice were challenged intranasally with 1 × 10⁶ PFU VACV-WR at Day 150 following a single prime immunization. This challenge dose maintained complete lethality in unvaccinated animals while providing sufficient resolution to distinguish protective efficacy among vaccination regimens at an extended durability time point.

Unvaccinated animals developed progressive weight loss beginning approximately 3–4 dpi and succumbed uniformly by 7 dpi (Fig. 7A, B). Mice receiving a single dose of parental MVA exhibited transient morbidity and partial protection, with one mortality observed following infection. In contrast, both MVA-X Prime and MVA Prime/Boost vaccination maintained stable body weight and complete survival throughout the observation period, demonstrating sustained protective immunity at an extended post-vaccination time point.

**Figure 7:**
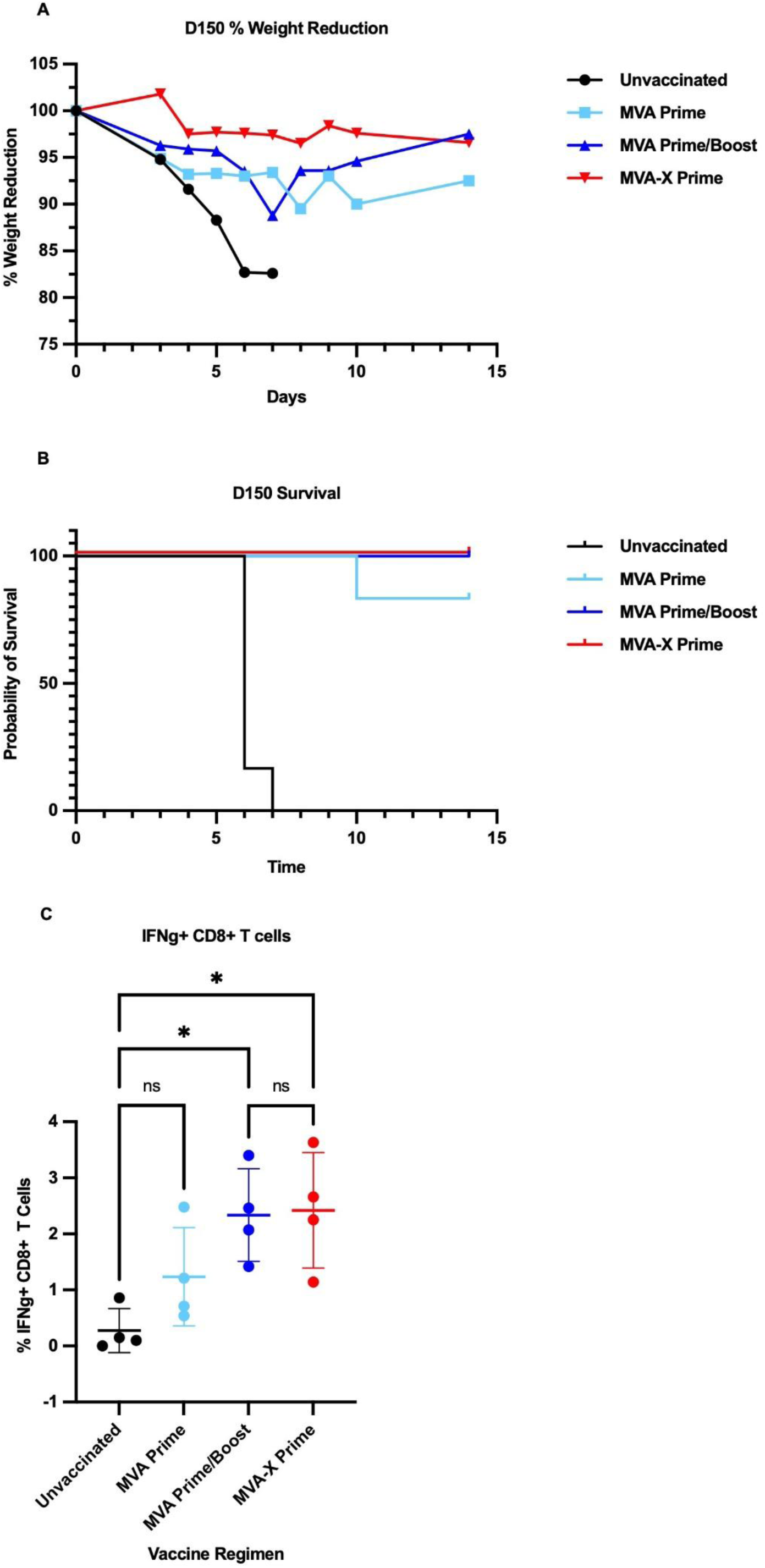
MVA-X Prime maintains long-term protection and durable CD8⁺ T cell responses at Day 150 post-prime vaccination. Mice were challenged intranasally with 1x 10^6^ PFU VACV-WR at Day 150 post-prime vaccination and monitored for disease progression and survival. Intracellular cytokine staining was performed using a separate cohort of vaccinated mice at Day 150 following ex vivo MVA stimulation. Individual points represent individual animals (n = 5–6 per group); horizontal bars indicate mean ± s.d. **(A)** Percent body weight relative to pre- challenge baseline over 14 d post-infection (dpi). Unvaccinated animals developed progressive weight loss beginning at approximately 3–4 dpi and reached humane endpoint criteria by 7 dpi. MVA-X Prime and MVA Prime/Boost groups maintained stable body weights throughout the observation period, whereas the Single-dose parental MVA group exhibited moderate transient weight loss with partial recovery. Group averages are shown. **(B)** Kaplan–Meier survival analysis following Day 150 challenge. All unvaccinated animals succumbed to infection by 7 dpi, with five deaths occurring at 6 dpi and one additional death at 7 dpi. Complete survival was maintained in the MVA-X Prime and MVA Prime/Boost groups throughout the study period, while a single mortality occurred in the MVA Prime group at 10 dpi, resulting in partial protection. Survival curves were analyzed using the log-rank (Mantel–Cox) test. **(C)** Percentage of IFNγ⁺ CD8⁺ T cells following ex vivo MVA stimulation at Day 150 post-prime vaccination. Cytokine-producing CD8⁺ T cells were identified following sequential gating on singlets, viable lymphocytes, CD3⁺ T cells, and CD8⁺ T cells. Both MVA-X Prime and MVA Prime/Boost groups demonstrated elevated IFNγ-producing CD8⁺ T cell responses relative to unvaccinated controls, while the Single-dose parental MVA group showed an intermediate response profile. Statistical analysis was performed using ordinary one-way ANOVA with Tukey’s multiple-comparisons test. Exact statistical comparisons are indicated in the figure. ns, not significant; *P < 0.05.

To determine whether durable protection remained associated with antigen-specific cellular immunity, intracellular cytokine staining was performed using a separate cohort of vaccinated animals at Day 150 post-prime vaccination. Following ex vivo stimulation, both MVA-X Prime and MVA Prime/Boost groups exhibited increased frequencies of IFNγ-producing CD8⁺ T cells relative to unvaccinated controls, whereas the MVA Prime group displayed an intermediate response profile (Fig. 7C). These findings demonstrate that a single MVA-X vaccination supports durable antigen-specific CD8⁺ T-cell responses that persist through long-term protective immunity.

### MVA-X vaccination preserves protection against Clade I MPXV challenge in susceptible CAST/EiJ mice model

To determine whether the protective efficacy of MVA-X extended beyond VACV-WR, vaccinated CAST/EiJ mice were challenged intranasally with a lethal dose (1×10^6^ PFU) of Clade I MPXV (Zaire strain) at Day 55 post-prime vaccination. This highly susceptible mouse strain provides a stringent model for evaluating vaccine efficacy against severe Mpox disease.

Unvaccinated animals developed pronounced weight loss beginning at 5 dpi, consistent with progressive disease following Clade I MPXV infection (Fig. 8A). In contrast, single-dose MVA-X vaccination-maintained body weight near baseline throughout infection, indicative of minimal clinical disease. Although MVA Prime/Boost vaccination also protected against mortality, animals in this group exhibited greater morbidity, characterized by increased weight loss and substantial inter-animal variability.

**Figure 8:**
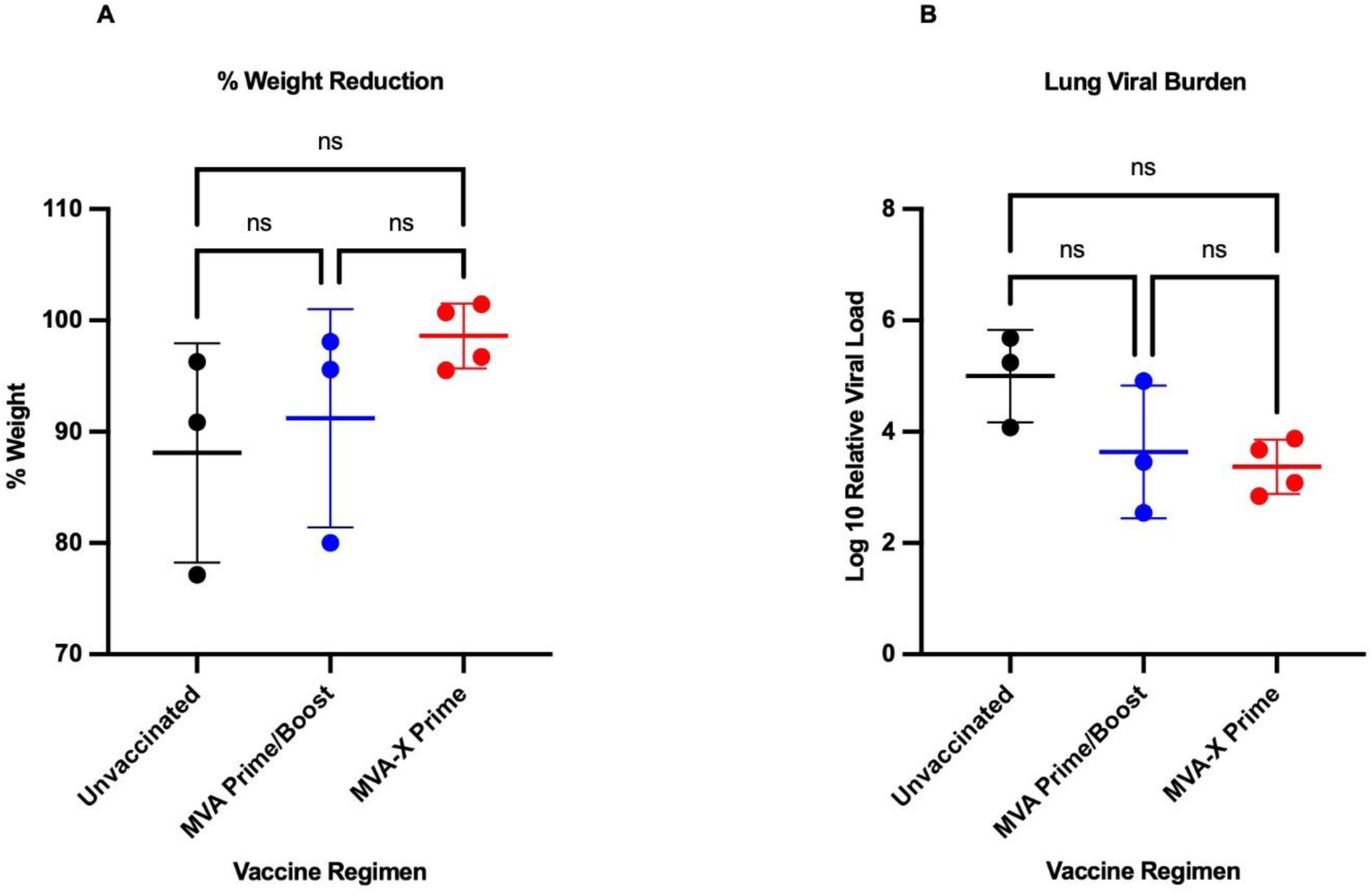
MVA-X Prime vaccination preserves body weight following Clade I MPXV challenge in CAST/EiJ mice. **(A)** CAST/EiJ mice were challenged with Clade I MPXV and monitored for disease progression. Percent body weight relative to pre-infection baseline was measured at 5 d post-infection (dpi). Individual points represent individual animals; horizontal bars indicate mean ± s.d. Mice vaccinated with MVA-X demonstrated improved weight maintenance compared with formulation buffer controls, whereas the parental MVA group showed greater inter-animal variability in disease outcome. Statistical analysis was performed using ordinary one-way ANOVA with Tukey’s multiple-comparisons test. Exact statistical comparisons are indicated in the figure. ns, not significant; *P < 0.05. **(B)** Relative pulmonary viral genome levels measured in lung tissue collected at 5 d post-infection (dpi) following Day 55 intranasal challenge with 1 × 10^6^ PFU MPXV Clade I. Viral DNA was quantified by qPCR targeting the VACV A12L gene and normalized to β-actin expression. Relative viral expression is presented as log₁₀(2^−ΔΔCt) + 5, where the unvaccinated group served as the reference comparator for ΔΔCt calculations and 5 was added as a visualization constant. Individual points represent individual animals (n = 4 per group); horizontal bars indicate mean ± s.d. Unvaccinated group exhibited high pulmonary viral burden, whereas MVA-X Prime animals exhibited a trend toward reduced pulmonary viral genome burden relative to unvaccinated controls. MVA Prime/Boost group showed high inter-animal variability

Analysis of viral genome burden demonstrated a trend toward reduced tissue-associated viral levels in MVA-X-vaccinated animals compared with unvaccinated controls, although this difference did not reach statistical significance (p = 0.087; Fig. 8B). Together, these findings demonstrate that the protective effects of MVA-X extend beyond VACV-WR challenge and remain effective against highly pathogenic Clade I MPXV.

## Discussion

A central challenge in vaccine development is achieving durable protection after a single immunization while preserving an established safety profile. Here, we demonstrate that transient immune checkpoint modulation embedded directly within an attenuated MVA vector enhances long-term protective immunity without altering vector attenuation or requiring booster immunization. A single dose of MVA-X conferred protection comparable to a conventional two-dose MVA regimen across multiple orthopoxvirus challenge models supporting the concept that improving early immune priming can generate durable protection comparable to conventional prime-boost vaccination. Together, these findings support the concept that early immune programming, rather than sustained humoral immunity alone, can establish durable vaccine-mediated protection.

A notable observation supporting this hypothesis was the discordance between humoral immunity and protective efficacy. Although both binding and neutralizing antibody responses progressively declined following vaccination, MVA-X-vaccinated animals remained completely protected against lethal VACV challenge through Day 150. These findings indicate that maintenance of high circulating antibody titers was not essential for long-term protection in this vaccine model and suggest that additional immune mechanisms sustain antiviral immunity after antibody responses contract. Although neutralizing antibodies remain important correlates of protection for many viral vaccines, they may not fully capture the durability of protection elicited by vaccine platforms that generate robust cellular immune responses^47–50^.

Consistent with this interpretation, pre-challenge depletion of CD8⁺ T cells resulted in a marked increase in viral burden in MVA-X-vaccinated animals, demonstrating a substantial contribution of vaccine-induced CD8⁺ T-cell immunity to viral control following lethal orthopoxvirus challenge. Collectively, these findings support functional CD8⁺ T-cell responses as an important correlate of durable orthopoxvirus immunity, particularly after circulating antibody responses have contracted.

Consistent with this interpretation, MVA-X elicited sustained antigen-specific CD8⁺ T-cell responses characterized by increased frequencies of IFNγ⁺ and TNFα⁺ CD8⁺ T cells. The persistence of IFNγ-producing CD8⁺ T cells through Day 150 is particularly notable given the established role of IFNγ in restricting orthopoxvirus replication, enhancing antigen presentation, and promoting viral clearance^26^. Increased TNFα-producing CD8⁺ T cells at Day 90 further supports enhanced effector function, as TNFα acts synergistically with IFNγ to coordinate antiviral immune responses and improve control of infected tissues^51^. In contrast, IL-2⁺ CD8⁺ T cell responses were less distinctly separated across vaccination groups, suggesting preferential enhancement of effector-associated cytokine responses rather than broad expansion of all functional subsets. Importantly, pre-challenge CD8⁺ T cell depletion resulted in increased viral burden in MVA-X-vaccinated animals, supporting a functional role for vaccine-induced CD8⁺ T cell immunity in mediating viral control following lethal orthopoxvirus challenge. This is particularly important because CD8⁺ T cells play a central role in orthopoxvirus clearance, whereas attenuated viral vectors such as MVA often induce comparatively limited T cell priming following a single-dose^52^. In contrast, NK cell depletion had minimal effect on pulmonary viral burden, indicating that the enhanced protection conferred by MVA-X is predominantly mediated through adaptive CD8⁺ T-cell immunity rather than NK cell-dependent antiviral responses.

The enhanced cellular immunity observed following MVA-X vaccination was accompanied by marked suppression of viral replication at the primary site of infection. Both pulmonary viral genome levels and infectious viral titers were substantially reduced following VACV challenge, while histopathological analysis demonstrated preservation of lung architecture despite evidence of active immune infiltration. These findings suggest that MVA-X-mediated immunity acts early after infection to restrict viral replication before extensive tissue damage develops. Importantly, preservation of pulmonary architecture in the presence of organized immune infiltrates indicates that protection was associated with effective antiviral immune activity rather than an absence of inflammation. This distinction is particularly relevant in orthopoxvirus disease, where pathology reflects both direct viral replication and the inflammatory consequences of uncontrolled infection^53^. Together, the coordinated reduction in pulmonary viral burden, systemic dissemination, and tissue pathology indicates that MVA-X improves the quality of protection by promoting earlier containment of infection and limiting subsequent disease progression.

The design of MVA-X was intended to provide localized and transient PD-1 antagonism during the critical window of vaccine-induced immune priming while avoiding the risks associated with systemic checkpoint blockade. Following vaccination, MVA preferentially infects professional antigen-presenting cells, including dendritic cells, which migrate to draining lymphoid tissues and initiate antigen-specific T-cell activation. Within the MVA-X platform, infected antigen-presenting cells express and secrete the LD10 peptide, enabling localized antagonism of PD-1 signaling during antigen presentation. Because MVA is replication-deficient in mammalian cells and LD10 exhibits a short in vivo half-life, checkpoint modulation is expected to be restricted both spatially and temporally to the early stages of the immune response^37^.

This targeted delivery strategy is proposed to transiently relieve PD-1-mediated inhibition at the site of antigen presentation while minimizing the prolonged systemic immune activation associated with conventional checkpoint inhibitors^54–56^. The persistence of antigen-specific IFNγ-producing CD8⁺ T cells through Day 150, together with the increased frequency of effector memory CD8⁺ T cells observed at Day 90, suggests that transient PD-1 modulation during vaccination influences not only effector function but also the establishment of long-lived cellular immunity. Effector memory T cells mediate rapid recall responses following pathogen re-exposure and have been associated with improved control of orthopoxvirus infection. Although additional studies are needed to define the mechanisms governing memory differentiation, these findings support the hypothesis that localized PD-1 antagonism during immune priming establishes durable cellular immunity by shaping early immune programming events that persist long after checkpoint modulation has ceased. Importantly, this strategy preserves the established safety profile of the MVA platform while enhancing the durability and functional quality of vaccine-induced cellular immunity following a single immunization^7,9,52,57^.

Importantly, the protective effects of MVA-X were not restricted to the VACV-WR challenge model. In the highly susceptible CAST/EiJ mouse model, single-dose MVA-X vaccination protected against lethal Clade I MPXV challenge, supporting the broader applicability of this approach across distinct orthopoxviruses. The inclusion of Clade I MPXV is particularly relevant because these viruses are associated with greater virulence, higher case fatality rates, and more severe clinical disease than the Clade II viruses responsible for the 2022 global outbreak^58–60^. Human Mpox is characterized by febrile illness, lymphadenopathy, and progressive mucocutaneous lesions that may persist for weeks and, in severe cases, progress to secondary bacterial infections, pneumonitis, encephalitis, ocular complications, and death^53,61^. Consequently, protection against Clade I MPXV provides a clinically relevant assessment of vaccine performance against severe orthopoxvirus disease. Although both MVA-X and MVA Prime/Boost protected against mortality following Clade I MPXV challenge, MVA-X-vaccinated animals exhibited more consistent maintenance of body weight and a trend toward reduced tissue-associated viral burden. Because body weight loss is a sensitive indicator of disease severity in orthopoxvirus infection, these findings suggest that MVA-X not only prevents mortality but also improves control of disease following exposure. While the reduction in viral genome burden did not reach statistical significance, the observed trend, together with the improved clinical outcomes, supports the conclusion that checkpoint- integrated MVA vaccination retains protective efficacy against a highly pathogenic MPXV challenge model. Collectively, these findings extend the relevance of the MVA-X platform beyond vaccinia-based challenge systems and support its broader potential for protection against clinically important orthopoxvirus infections.

The ability of MVA-X to confer durable protection after a single immunization has important implications for outbreak preparedness, where rapid vaccine deployment and incomplete booster compliance can limit the effectiveness of multidose vaccination strategies. MVA-based vaccines are well suited for large-scale manufacturing, lyophilization, and stockpiling, and improving their single-dose efficacy could substantially increase their operational flexibility during emergency responses. More broadly, incorporating transient immune checkpoint modulation directly within the vaccine vector represents a modular strategy that may be adaptable to additional vaccine platforms and pathogens for which durable cellular immunity is critical for protection^48,62,63^.

Several limitations should be considered when interpreting these findings. Although our data demonstrate a substantial contribution of CD8⁺ T cells to MVA-X-mediated protection, this study did not directly define the molecular mechanisms by which transient PD-1 antagonism influences T-cell differentiation or memory formation. Likewise, all memory subset composition, tissue-resident T-cell responses, and markers of activation or exhaustion were not examined and warrant further investigation. In addition, while the VACV-WR challenge model provides a rigorous and well-established platform for evaluating orthopoxvirus vaccine efficacy, future studies will be required to determine whether this strategy provides similar benefits across additional antigens, viral vectors, and disease settings. Defining how checkpoint-integrated vaccination shapes long-term immune memory, recall responses, and tissue-specific immunity will further clarify the broader applicability of this platform.

In summary, transient immune checkpoint modulation incorporated directly within an attenuated MVA vector enhanced the magnitude and durability of protective immunity following a single immunization. Despite progressive contraction of circulating antibody responses, MVA-X maintained long-term protection against lethal VACV challenge, reduced viral replication and pulmonary pathology, elicited durable antigen-specific CD8⁺ T-cell responses, and retained efficacy against highly pathogenic Clade I MPXV. Together, these findings demonstrate that localized, vaccine-intrinsic checkpoint modulation can improve the functional durability of established viral-vector vaccines while preserving their favorable safety characteristics. More broadly, this work establishes a modular strategy for enhancing single-dose vaccine efficacy and supports the development of next-generation vaccines capable of providing durable protection against emerging infectious diseases.

## Methods

### Ethics Statement

All animal studies were conducted in accordance with the Guide for the Care and Use of Laboratory Animals, the Public Health Service Policy on Humane Care and Use of Laboratory Animals and ARRIVE guidelines. Experimental protocols were approved by the Washington State University Institutional Animal Care and Use Committee. Washington State University is accredited by the Association for Assessment and Accreditation of Laboratory Animal Care International (AAALAC-000480). Animals were housed in HEPA-filtered individually ventilated cages with ad libitum access to food and water and monitored daily by trained personnel. Mice were euthanized at predefined humane endpoints or terminal study time points by ketamine–xylazine overdose (9 mg/kg ketamine, 1 mg/kg xylazine) followed by bilateral thoracotomy.

### Vaccine construction and viral propagation

A recombinant modified vaccinia Ankara (MVA) vector expressing the peptide-based PD-1 antagonist LD10 (MVA-X) was generated as previously described^37^. Briefly, five tandem copies of the LD10 coding sequence, each preceded by a tissue plasminogen activator (tPA) secretion signal peptide and separated by proteolytic cleavage motifs, were inserted into the MVA genome under the control of an early/late promoter to facilitate peptide secretion. Recombinant viruses were generated by homologous recombination, propagated in DF-1 cells, and titrated by standard plaque assay on Vero-81 cells.

### Animals and vaccination regimen

Eight- to ten-week-old age- and sex-matched C57BL/6J mice (The Jackson Laboratory) were maintained under specific pathogen-free conditions. Unless otherwise indicated, experiments included six mice per group. Mice were anesthetized with a ketamine–xylazine cocktail and immunized intramuscularly with 5 × 10⁶ PFU of vaccine in formulation buffer. Vaccination groups included formulation buffer control (FB/FB), single-dose parental MVA (MVA Prime), two-dose parental MVA (MVA Prime/Boost), and single-dose MVA-X (MVA-X Prime). FB/FB mice received formulation buffer on Days 0 and 28; MVA Prime mice received formulation buffer on Day 0 and parental MVA on Day 28; MVA Prime/Boost mice received parental MVA on Days 0 and 28; and MVA-X Prime mice received formulation buffer on Day 0 and MVA-X on Day 28. Single-dose regimens were administered on Day 28 to align with the final immunization of the MVA Prime/Boost group, enabling comparison of immune responses and protective efficacy at equivalent intervals after the most recent vaccine exposure. Blood samples were collected longitudinally via the saphenous vein for serological analyses.

### VACV challenge models

Protective efficacy was evaluated using intranasal challenge with Vaccinia virus Western Reserve (VACV-WR). For high-dose challenge studies, mice were challenged with 1 × 10⁷ PFU VACV-WR on either Day 55 or Day 90 post-prime vaccination. To assess the long-term durability of protection, a separate cohort was challenged with 1 × 10⁶ PFU VACV-WR on Day 150 post-prime vaccination. This challenge dose was selected because it maintained lethality in unvaccinated controls while providing greater resolution of protective differences among vaccinated groups during extended durability studies. VACV-WR stocks were propagated and titrated on Vero-81 cells prior to challenge. Mice were anesthetized with a ketamine–xylazine cocktail and inoculated intranasally with 5 μL of virus suspension into the left nare. Animals were monitored daily for body weight, clinical signs of disease, and survival for 14 days following infection. Humane endpoint criteria included greater than 20% body weight loss or clinical signs consistent with severe disease, at which point animals were euthanized according to approved IACUC protocols.

### Sample collection and processing

Blood samples were collected longitudinally via the saphenous vein at designated time points for serological analyses. For tissue-based studies, subsets of mice were euthanized at specified time points following VACV-WR or MPXV challenge, and whole blood was collected by cardiac puncture for serum isolation. Lungs, spleens and livers were collected for virological, immunological, and histopathological analyses. Tissues were divided at necropsy, with one portion fixed in 10% neutral buffered formalin for histopathological evaluation and the remaining portion snap-frozen and stored at −80°C for downstream analyses, including viral burden quantification. All samples were processed and stored under appropriate conditions until analysis.

### Binding antibody ELISA

MVA-specific IgG responses were quantified by enzyme-linked immunosorbent assay (ELISA). High-binding 96-well plates (Thermo Scientific MaxiSorp) were coated overnight at 4°C with 1 × 10⁶ PFU purified MVA/well diluted in phosphate-buffered saline (PBS). Plates were blocked with StartingBlock™ Blocking Buffer (Thermo Scientific) prior to incubation with serially diluted mouse plasma samples. Plasma samples were initially diluted 1:320 and subjected to seven consecutive two-fold serial dilutions. Following washing with PBS containing 0.05% Tween-20 (PBS-T), plates were incubated with horseradish peroxidase-conjugated goat anti-mouse IgG secondary antibody (VWR; 1:4,000 dilution). Bound antibodies were detected using SureBlue™ TMB substrate (SeraCare), and the reaction was terminated with 1 N HCl. Absorbance was measured at 450 nm using a Varioskan™ LUX microplate reader. Endpoint titers were defined as the reciprocal of the highest plasma dilution producing an absorbance value greater than two-fold above background. Endpoint titers were used for all statistical analyses.

### Fluorescent Neutralization Assay D90 Post Prime

Neutralizing antibody activity was assessed using a fluorescent VACV-E3L-GFP reporter virus. Vero-81 cells were seeded in 96-well tissue culture plates at 5 × 10⁴ cells per well one day prior to infection. Aliquots of serum samples from each animal were heat-inactivated at 56 °C for 30 min and serially diluted in 2% MEM medium. The multiplicity of infection required to achieve approximately 80% GFP-positive Vero-81 cells was empirically determined before the assay. Diluted serum samples were incubated with VACV-E3L-GFP for 2 h at 37 °C before addition to Vero-81 monolayers. Virus-serum mixtures were then incubated with cells for 8 h at 37 °C in cell culture incubator.

Following infection, medium was removed and cells were washed once with PBS. Cells were detached with 30 μL trypsin-EDTA per well and incubated at 37 °C until the monolayer loosened for 2-3 minutes. Trypsin was neutralized by adding 90 μL of 2% serum-containing medium. Cells were transferred to U-bottom 96-well plates and centrifuged at 400 × g for 4 min at 4 °C. Supernatants were removed, and cells were washed once with PBS prior to fixation with 4% paraformaldehyde for 15 min at room temperature. Fixed cells were washed once and resuspended in FACS buffer for acquisition.

Samples were acquired on a Cytek Aurora spectral flow cytometer. GFP-positive infected cells were detected using the Alexa Fluor 488 channel. Analysis was performed by sequential gating on forward and side scatter to exclude debris, singlet discrimination, and identification of GFP-positive virus-infected cells (Supplemental Figure 2).

Percent neutralization was calculated relative to virus-only wells at the same MOI:

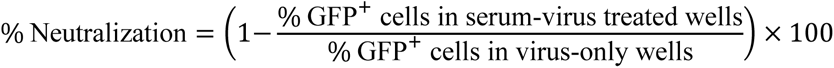

### Viral titers and quantitative PCR

Tissues were collected at euthanasia (5 dpi) from the Day 90 challenge cohort representing the four vaccination groups (Unvaccinated, MVA Prime, MVA Prime/Boost, and MVA-X Prime). Lung tissues were homogenized at a ratio of 10 mg tissue per 100 μL RPMI-based medium. Samples underwent two freeze–thaw cycles before mechanical homogenization. An aliquot (100 μL) was reserved for DNA extraction and quantitative PCR (qPCR), while the remaining homogenate underwent an additional freeze–thaw cycle followed by sonication (20% amplitude, 1 min) to maximize viral release.

Vero-81 cells were seeded in 6-well plates at a density of 4 × 10⁵ cells/well one day before infection. Serial 10-fold dilutions (10⁻¹–10⁻⁶) of tissue homogenates were prepared in 2% MEM, and 100 μL of each dilution was added to duplicate wells. Following a 1 h adsorption at 37°C with gentle rocking every 10 min, cells were washed with PBS, overlaid with 2 mL of 2% MEM, and incubated for 36–48 h. Monolayers were fixed and stained with crystal violet for at least 1 h, rinsed with deionized water, air dried, and plaques were enumerated independently by two blinded scorers. Viral titers were expressed as PFU/g of tissue.

DNA was extracted from lung tissue homogenates and EDTA-treated whole blood using the Monarch® Genomic DNA Purification Kit (New England Biolabs, Cat#T3010) according to the manufacturer’s instructions. DNA was quantified before storage at −20°C. Vaccinia virus genomes were quantified by qPCR targeting the A12L gene using 2× PrimeTime® Gene Expression Master Mix (IDT) on a QuantStudio™ 6 Pro Real-Time PCR System (Applied Biosystems). Reactions (20 μL) were performed in duplicate using 2 μL of tissue DNA or 5 μL of blood DNA. Thermal cycling consisted of 50°C for 2 min, 95°C for 3 min, and 40 cycles of 95°C for 15 s, 50°C for 15 s, and 60°C for 1 min. Viral genome levels were normalized to β- actin using the 2^−ΔΔCq method with the unvaccinated group as the calibrator. Values were transformed as log₁₀(2^−ΔΔCq) + 5 for graphical visualization, and statistical analyses were performed using one-way ANOVA with Tukey’s multiple-comparison test in GraphPad Prism.

### Histopathology

Lung tissues were fixed overnight in 10% neutral-buffered formalin, transferred to 70% ethanol, and processed through graded ethanol, xylene, and paraffin using standard histological procedures before paraffin embedding. Tissue blocks were sectioned at 5 μm for hematoxylin and eosin (H&E) staining and immunohistochemistry.

For H&E staining, paraffin sections were deparaffinized in xylene, rehydrated through graded ethanol, and stained with hematoxylin followed by eosin using standard protocols. Sections were subsequently dehydrated, cleared in xylene, mounted with permanent mounting medium, and dried overnight before microscopic evaluation.

### Intracellular cytokine staining D90 Post Prime

To assess antigen-specific cellular immune responses, splenocytes were harvested at Day 90 post-prime vaccination and processed into single-cell suspensions following standard protocols^64^. Cells were seeded into V-bottom 96-well plates at a density of 2 × 10⁶ cells per sample and stimulated ex vivo with whole MVA at a multiplicity of infection (MOI) of 5. Two hours after stimulation, GolgiPlug (BD Biosciences) was added at a 1:1000 dilution.

For Fc receptor blockade, cells were incubated with anti-mouse CD16/32 Fc Block (BD Biosciences, Cat#553142) diluted 1:100 in FACS buffer for 30 min on ice. Viability staining was performed using Fixable Viability Stain 700 (FVS700; BD Biosciences, Cat#564997) prepared at 3 μL per 10 mL staining buffer, with 100 μL added per sample for 20 min in the dark.Surface staining was performed using antibodies against CD45 (FITC; BioLegend, Cat#109806), CD19 (eFluor 450; Invitrogen, Ref#48-0193-82), CD3 (BUV737; BD Biosciences, Cat#564380), CD8α (BV510; BD Biosciences, Cat#563068), and CD4 (BUV496; BD Biosciences, Cat#564667) prepared in FACS buffer. Cells were incubated with 100 μL antibody cocktail for 20 min in the dark.

Cells were fixed and permeabilized using the BD Cytofix/Cytoperm kit (BD Biosciences, Cat#554714) according to the manufacturer’s instructions before intracellular staining with antibodies against IFNγ (APC; Invitrogen/eBioscience, Ref#17-7311-82), TNFα (BV711; BD Biosciences, Cat#563944), and IL-2 (PE-CF594; BD Biosciences, Cat#562483). Cells were incubated with 100 μL intracellular staining cocktail for 20 min in the dark and resuspended in FACS buffer for acquisition.

Samples were acquired on a Cytek Aurora spectral flow cytometer within 24 h of staining. Spectral unmixing and compensation were performed using unstained controls, viability controls, and UltraComp eBeads (Invitrogen, Ref#01-2222-42) stained with individual antibodies. A minimum of 50,000 events were collected per sample. Sequential gating identified singlets, viable lymphocytes, CD3⁺ T cells, and CD8⁺ T-cell populations, followed by quantification of IFNγ⁺, TNFα⁺, and IL-2⁺ CD8⁺ T cells. Gates were established using unstimulated and healthy control samples (Supplementary Fig. 3).

### In Vivo CD8⁺ T cell depletion and challenge study

To determine the contribution of CD8⁺ T cells to vaccine-mediated protection, in vivo CD8⁺ T cell depletion was performed prior to lethal VACV-WR challenge. Mice were vaccinated with formulation buffer (FB/FB), homologous MVA-Prime/Boost, or Single-dose MVA-X-Prime according to the vaccination schedule described above. At Day 90 post-prime vaccination, animals received intraperitoneal injections of 250 μg anti-mouse CD8α depleting antibody (clone 2.43, Bio X Cell) on Days -3 and -1 prior to challenge. Control animals received matched isotype antibody using the same schedule and dosing regimen. Peripheral blood was collected prior to infection to confirm efficient depletion of circulating CD8⁺ T cells by flow cytometry (Supplemental Figure 4).

Following depletion, mice were challenged intranasally with 1 × 10⁷ PFU VACV-WR. Blood and tissues were collected at 5dpi for assessment of viral burden. Viral genome quantification was performed by qPCR targeting the VACV A12L gene following DNA extraction using the Monarch Genomic DNA Purification Kit. Relative viral levels were normalized to β-actin and calculated using the 2^−ΔΔCt method.

### Intracellular cytokine staining (ICS) assay at Day 150 post-prime vaccination

To assess long-term antigen-specific cellular immune responses, splenocytes were harvested at Day 150 post-prime vaccination and processed into single-cell suspensions following standard protocols. Cells were adjusted to a final concentration of 1 × 10⁶ cells per 100 μL and plated in 96-well plates for ex vivo stimulation.

Splenocytes were stimulated with parental MVA at a multiplicity of infection (MOI) of 1 PFU/cell. Unstimulated and Cell Stimulation Cocktail-treated cells served as negative and positive controls, respectively. Following a 2 h incubation at 37°C with 5% CO₂, GolgiPlug

(Brefeldin A; BD Biosciences, Cat#555029) and GolgiStop (Monensin; BD Biosciences, Cat#554724) were added at 40 μL/mL each, followed by an additional 5–6 h incubation.

Cells were stained with Viobility™ 405/520 Fixable Dye prepared by diluting 50 μL dye in 12.5 mL PBS, with 100 μL added per well for 15 min. Surface staining was performed using CD3ε (PerCP-Vio® 700, Miltenyi Biotec), CD4 (VioBlue®, Miltenyi Biotec), and CD8α (APC-Vio® 770, Miltenyi Biotec) antibodies prepared as a master mix containing 2 μL of each antibody in 94 μL flow staining buffer per sample and incubated for 15 min.

Cells were fixed and permeabilized using Cytofix/Cytoperm solution for 20 min at 2–8°C before intracellular staining with anti-mouse IFNγ-APC (0.75 μL/sample) diluted in Perm/Wash buffer to a final volume of 100 μL per well. Following a 15 min incubation, cells were washed, resuspended in PBS, and acquired on a flow cytometer. Data was analyzed using FlowJo software. Sequential gating identified singlets, viable lymphocytes, CD3⁺ T cells, and CD8⁺ T-cell populations before quantification of IFNγ-producing CD8⁺ T cells.

### Clade I MPXV challenge model in CAST/EiJ mice

Age- and sex-matched male and female CAST/EiJ mice (8–10 weeks old; n = 3 per group) were vaccinated prior to challenge. Vaccination groups included mock-vaccinated controls (FB/FB), two-dose parental MVA vaccination (MVA Prime/Boost), and Single-dose MVA-X vaccination (FB/MVA-X). All Clade I MPXV challenge studies were conducted under biosafety level 3 (BSL-3) containment conditions.

A lethal intranasal challenge with 1 × 10⁶ PFU of Clade I MPXV (Zaire-79; BEI Resources NR-2324, V79-I-005) was performed 55 d post-prime vaccination under anesthesia. Animals were monitored daily for clinical signs of disease and body weight loss following infection.

Body weight measurements were recorded and expressed as percentage of pre-infection baseline body weight.

For viral genome quantification, DNA was extracted from lung homogenates using the Monarch Genomic DNA Purification Kit according to the manufacturer’s instructions. Quantitative PCR targeting the conserved VACV A12L gene was performed, and viral genome levels were normalized to β-actin expression. Relative viral DNA abundance was calculated using the 2^−ΔΔCt method, with the mock-vaccinated group used as the reference comparator.

### Statistical analysis

Statistical analyses were performed using GraphPad Prism. Survival curves were analyzed using the log rank Mantel Cox test. Group comparisons were performed using one way ANOVA with appropriate multiple comparisons testing as indicated in the figure legends. Data are presented as mean with standard deviation unless otherwise stated. Statistical significance was defined as P less than 0.05.

## Authors Contribution

H.K. designed the study, performed experiments, and contributed to writing and editing the manuscript. R.M., J.D., B.G., S.B., B.O., B.A., and C.P.L. assisted with experimental assays. M.H, A.D., and S.RO. contributed to vaccine design and generation, assay development, experimental design, and manuscript editing. A.K. assisted in experimental design and execution, prepared the figures, and drafted the manuscript.

## Author contributions (Credit Taxonomy)

**Conceptualization:** H.K., A.K., M.H., A.D., S.R.O., M.N.

**Methodology:** H.K., A.K., M.H., S.R.O.

**Investigation:** H.K., A.K., R.M., J.D., C.P.L., M.H., S.R.O., B.G., S.B., B.O., B.A., P.K.

**Data curation:** H.K., A.K.

**Formal analysis:** H.K., A.K.

**Visualization:** A.K.

**Writing – original draft:** A.K., H.K.

**Writing – review & editing:** H.K., M.H., A.D., S.R.O., B.G., S.B.

**Supervision:** H.K.

## Data availability Statement

All data generated or analyzed during this study are included in this published article and its supplementary information files.

## Code availability

No custom code was used in this study.

## Declaration of generative AI use

The authors report that generative AI was not used in the study design, data collection, data analysis, interpretation of results, or generation of scientific conclusions. The authors used AI assistance via Grammarly for language refinement. All AI-assisted texts were reviewed and revised by the authors, who take full responsibility for the accuracy and integrity of the manuscript.

