## Supplemental Figures 1-3 for "Single-dose Efficacy of a Next-Generation Mpox Vaccine Harnessing an Immunomodulatory Peptide"

**Supplemental Figure 1:**

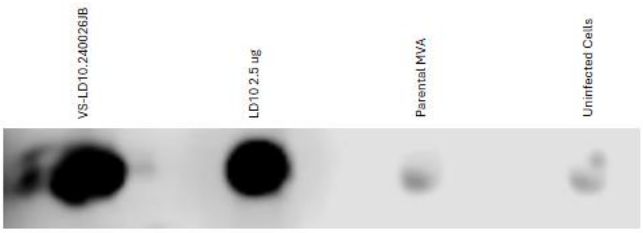


**Expression of LD10 by MVA-X infected cells**. DF-1 cells were infected with MVA-X, parental MVA, or mock infected. Forty-eight hours post infection, culture supernatants were collected and analyzed by dot blot alongside 2.5 μg of chemically synthesized LD10 peptide. Membranes were probed with an LD10-specific primary antibody followed by HRP-conjugated secondary antibody and visualized by chemiluminescence. A positive LD10-reactive signal was detected in supernatants from MVA-X-infected cells and the synthetic peptide control but not in parental MVA or mock-infected samples, confirming expression and secretion of LD10 by the recombinant vector.

**Supplemental Figure 2:**

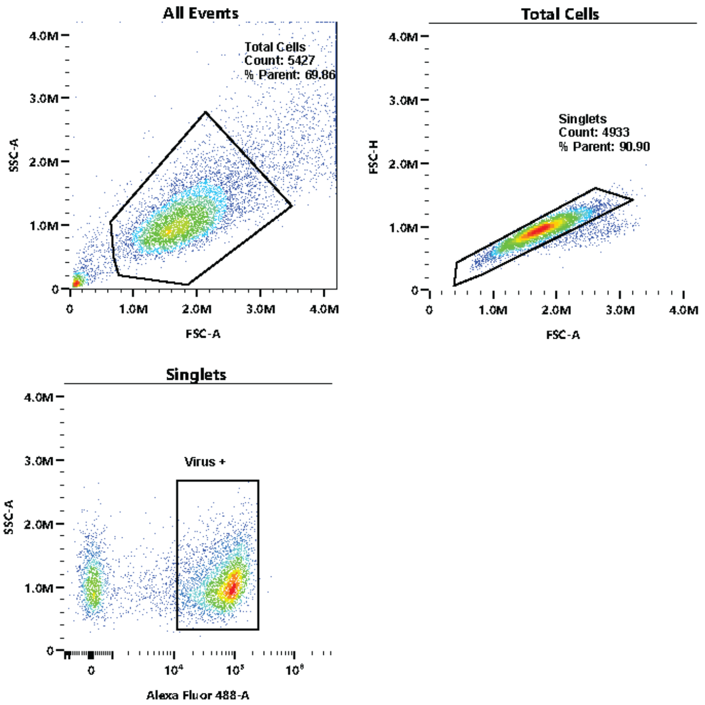


**Flow cytometry gating strategy for VACV-E3L-GFP neutralization assay.** Representative gating strategy used to quantify GFP-positive VACV-E3L-GFP infected Vero-81 cells during serum neutralization assays. Cells were first gated on forward and side scatter to exclude debris, followed by identification of singlets and quantification of GFP-positive infected cells. Neutralization was calculated as the reduction in GFP-positive cells relative to virus-only controls.

**Supplemental Figure 3:**

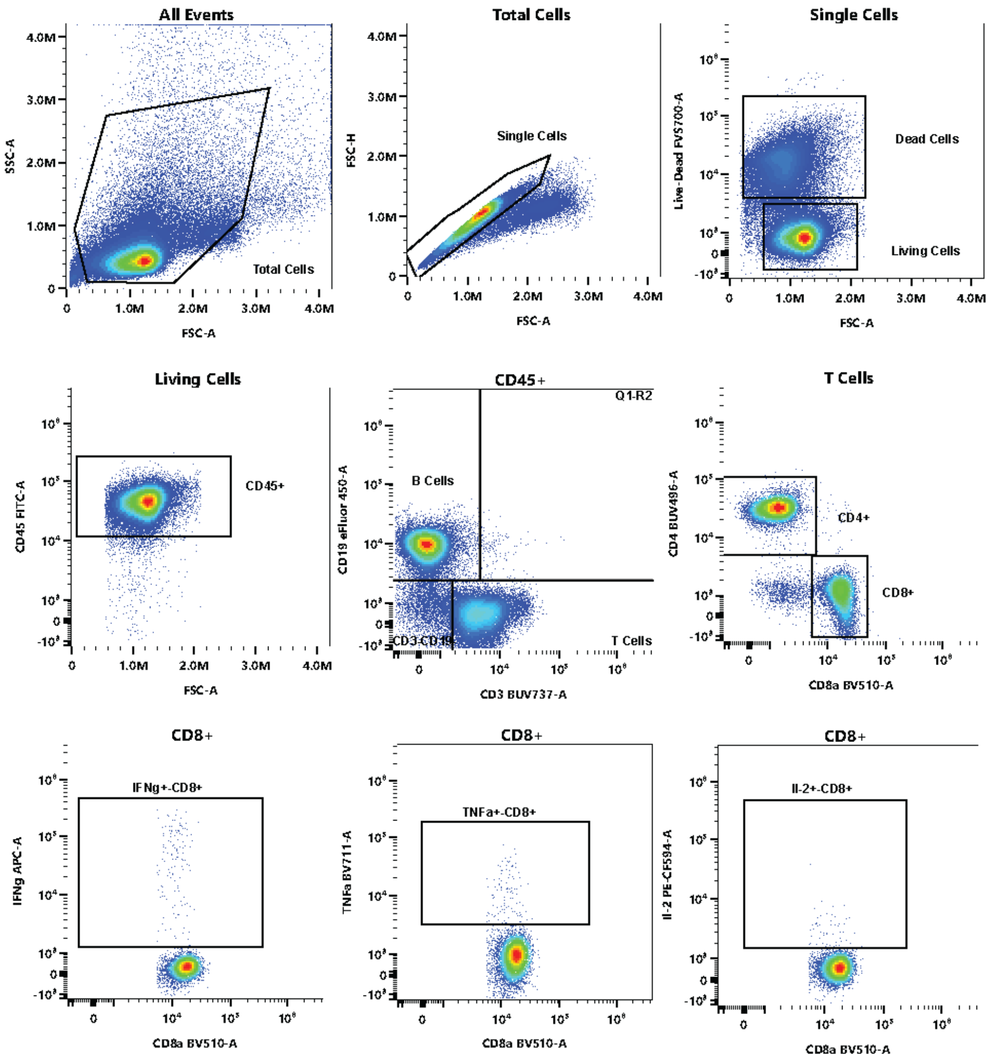


**Flow cytometry gating strategy for intracellular cytokine staining**. Representative gating strategy used for analysis of antigen-specific $CD8^+$ T-cell responses following ex vivo MVA stimulation. Sequential gates were applied to identify singlets, viable lymphocytes, $CD3^+$ T cells, and $CD8^+$ T cells, followed by quantification of $IFN\gamma^+$, $TNF\alpha^+$, and $IL\text{-}2^+$ cytokine-producing populations. Gates were established using unstimulated and healthy control samples.

**Supplemental Figure 4:**

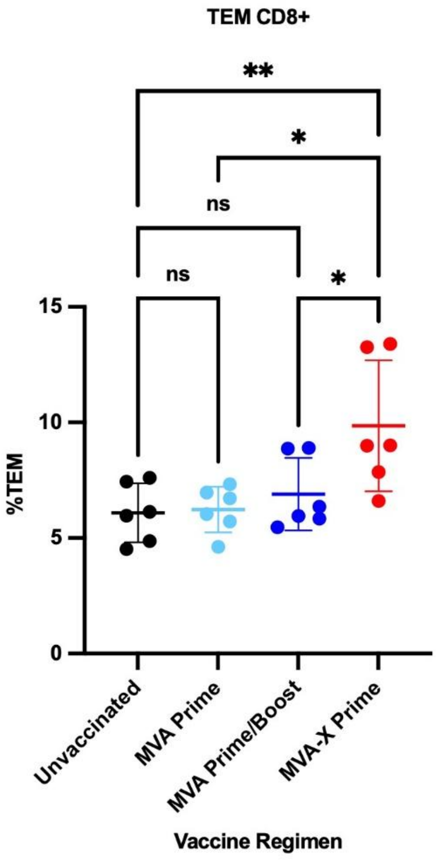


**MVA-X vaccination enhances effector memory CD8⁺ T-cell responses at Day 90 post-prime vaccination.** Splenocytes were collected from mice at Day 90 post-prime vaccination and analyzed by flow cytometry to quantify $CD8^+$ effector memory T (TEM; $CD44^+CD62L^-$) cells. MVA-X Prime vaccination resulted in significantly higher frequencies of $CD8^+$ TEM cells compared with unvaccinated, MVA Prime, and MVA Prime/Boost groups. Individual points represent individual mice (n = 5–6 per group), and horizontal bars indicate mean ± s.d. Statistical analysis was performed using ordinary one-way ANOVA with Tukey's multiple-comparisons test. Exact statistical comparisons are indicated in the figure. *$P < 0.05$; **$P < 0.01$.

**Supplemental Figure 5:**

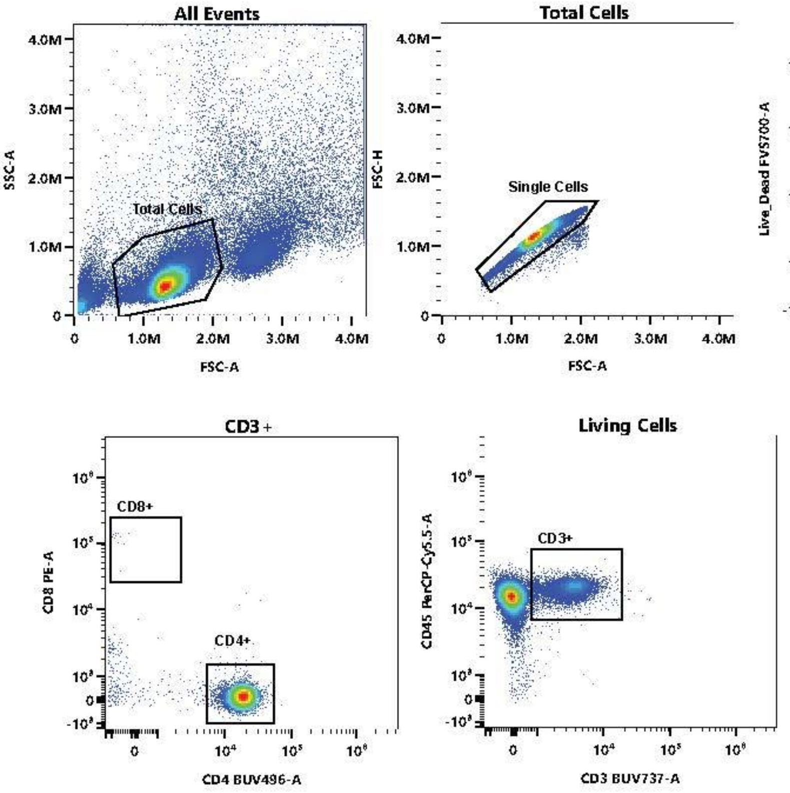


**Confirmation of in vivo CD8⁺ T-cell depletion**. Representative flow cytometry plots demonstrating depletion of circulating CD8⁺ T cells following administration of anti-CD8α antibody prior to VACV-WR challenge. Peripheral blood was collected immediately before infection and analyzed by flow cytometry. Efficient depletion of CD8⁺ T cells was confirmed in antibody-treated animals relative to undepleted controls.

**Supplementary Figure 6:**

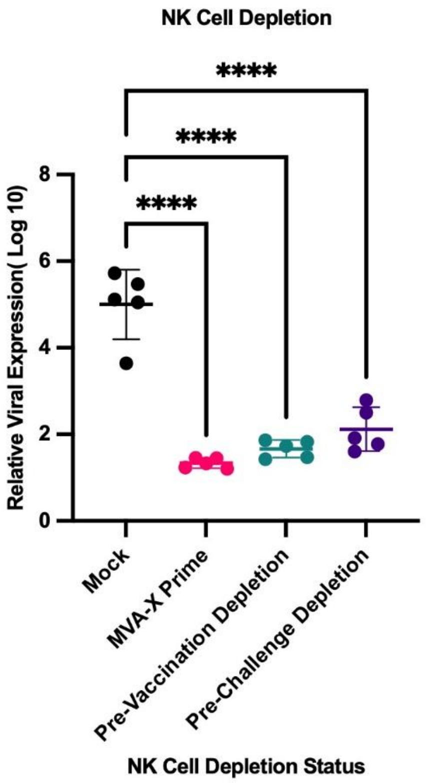


**Effect of NK cell depletion on pulmonary viral burden following lethal VACV-WR challenge:** Pulmonary viral burden in mock-vaccinated, non-depleted MVA-X-vaccinated, pre-vaccination NK cell-depleted, and pre-challenge NK cell-depleted mice following lethal VACV-WR challenge. Lung viral burden was quantified by qPCR at 5 dpi. Individual points represent individual mice (**n = 5** per group), and horizontal bars indicate mean ± s.d. Statistical analysis was performed using ordinary one-way ANOVA with Tukey's multiple-comparisons test. Exact statistical comparisons are indicated in the figure. ns, not significant.
